# SGLT2 inhibitors attenuate endothelial to mesenchymal transition and cardiac fibroblast activation

**DOI:** 10.1101/2023.12.18.572273

**Authors:** Kevin Schmidt, Arne Schmidt, Sonja Groß, Annette Just, Angelika Pfanne, Maximilian Fuchs, Maria Jordan, Elisa Mohr, Andreas Pich, Jan Fiedler, Thomas Thum

## Abstract

Beneficial effects of sodium glucose co-transporter 2 inhibitors (SGLT2is) in cardiovascular diseases have been extensively reported leading to the inclusion of these drugs in the treatment guidelines for heart failure. However, molecular actions especially on non-myocyte cells remain uncertain. We observed dose-dependent inhibitory effects of two SGLT2is, dapagliflozin (DAPA) and empagliflozin (EMPA), on inflammatory signaling in human umbilical vein endothelial cells (HUVECs). Proteomics analyses and subsequent enrichment analyses discovered profound effects of these SGLT2is on proteins involved in mitochondrial respiration and actin cytoskeleton. Validation in functional oxygen consumption measurements as well as tube formation and migration assays revealed strong impacts of DAPA. Considering that most influenced parameters played central roles in endothelial to mesenchymal transition (EndMT), we performed *in vitro* EndMT assays and identified substantial reduction of mesenchymal and fibrosis marker expression as well as changes in cellular morphology upon treatment with SGLT2is. In line, human cardiac fibroblasts (HCFs) exposed to DAPA showed less proliferation, reduced ATP production, and decelerated migration capacity while less extensive impacts were observed upon EMPA. Mechanistically, sodium proton exchanger 1 (NHE1) as well as sodium-myoinositol cotransporter (SMIT) and sodium-multivitamin cotransporter (SMVT) could be identified as relevant targets of SGLT2is in non-myocyte cardiovascular cells as validated by individual siRNA-knockdown experiments. In summary, we found comprehensive beneficial effects of SGLT2is on human endothelial cells and cardiac fibroblasts. The results of this study therefore support a distinct effect of selected SGLT2i on non-myocyte cardiovascular cells and grant further insights into potential molecular mode of action of these drugs.

## Introduction

Gliflozins are a class of small molecules derived from the molecular structure of the polyphenol phlorizin ^1^. They are primarily known for inhibiting the sodium-glucose cotransporter (SGLT) 2, which is expressed in epithelial cells in the proximal tubules of nephrons and responsible for reabsorption of 90% of the filtered glucose ^2^. Multiple clinical studies have demonstrated that the promotion of glucosuria by SGLT2 inhibitors (SGLT2is) can substantially alleviate diabetic burden ^3–6^ ensuing in their approval for the treatment of type 2 diabetes mellitus (T2D) ^7^. Interestingly, cardiovascular complications, a frequent comorbidity in T2D patients, were additionally reduced upon reception of SGLT2is ^4^. Further trials focusing on the impact of SGLT2is on heart failure (HF) with reduced ejection fraction independent of diabetic preconditions confirmed the cardiovascular benefits of SGLT2is ^8,9^. Recent trials also reported positive outcomes of dapagliflozin (DAPA) and empagliflozin (EMPA) in HF patients with preserved ejection fraction ^10,11^. Although apparent in large patient cohorts, the underlying mechanisms of cardiovascular effects of SGLT2is have still not been conclusively revealed, narrowing the imaginable spectrum of clinical application.

To some degree, the cardiovascular benefit could be attributed to the effect of SGLT2is in the kidney. The hindered reabsorption of glucose is disputed to be altering metabolic activity on an organismal level reducing glucose and free fatty acid oxidation while enhancing ketone body utilization in the failing heart thereby improving myocardial contractility and decelerating HF progression ^12,13^. Contrarily, other glucosuria promoting medications did not show such cardiovascular benefits ^14^. Therefore, alternative hypotheses have been proposed that SGLT2is rather mimic a nutrient deprivation condition than induce a shift to a different “fuel” ^15^. Thereby, low energy states instigate autophagic processes via sirtuin 1 and 5’ adenosine monophosphate-activated protein kinase signaling which, in turn, ultimately impact inflammatory processes and reactive oxygen species (ROS) accumulation ^14^.

However, these proposed systemic mechanisms cannot fully explain the cardiovascular-protective function of SGLT2is. While the complete cardiovascular system is lacking expression of SGLTs, existence of other targets of SGLT2is is probable and has been explored in recent years ^16^. As such, direct effects in the cardiovascular environment have been reported on interference with ion homeostasis in cardiomyocytes. Direct binding and inhibition of the sodium proton exchanger 1 (NHE1) have been shown in ammonium pulse assays ^17^, in which some groups also included cariporide as a positive control ^18^. Alteration of intracellular Na^+^ controls Ca^2+^ levels thereby improving contractility, regulating mitochondrial function and reducing ROS production ^19^. Impacts of SGLT2i seen in endothelial cells (ECs) are also disputed to be mediated through NHE1. While reported results have not consistently been reproducible, it has been proposed that inhibition of NHE1 by SGLT2i ultimately results in reduced ROS stress ^20,21^ and possibly a change in expression of adhesion molecules as well as interleukins (ILs) ^22,23^. Thereby, inflammatory processes are muted reducing overall stress factors and triggers of cardiac remodeling and vascular dysfunction in HF patients. Inflammatory signaling, in terms of NLR family, pyrin domain-containing 3 (NLRP3) abundance and pro-inflammatory cytokine expression, has also been reported to be reduced in cardiac fibroblasts (CFs) ^24^. Another study suggested that not only inflammation but rather general activation and myofibroblast differentiation of CFs is attenuated by SGLT2i ^25^. However, precise mechanistic insights to these observations are still elusive.

The aim of this study was to further define direct actions of SGLT2is in non-myocyte cardiovascular cells and to unravel molecular key nodes underlying observed effects. We show an impact of DAPA on HUVEC metabolism and angiogenic capacity and connect this to a modulation of mesenchymal activation of ECs as well as a reduction on fibroblast proliferation and migration. Based on siRNA-mediated gene silencing experiments of known and potential novel targets, we speculate on mechanisms of SGLT2is in considered cell types.

## Material and Methods

### General cell culture

Human embryonal kidney (HEK) 293FT received Dulbecco’s modified eagle’s medium, high glucose, (DMEM, 11965, Thermo Fisher Scientific, Waltham, MA, USA) containing 10% (volume/volume [v/v]) fetal bovine serum (FBS, 10270, Thermo Fisher Scientific), 1% (v/v) Penicillin-Streptomycin (PenStrep, 15070, Thermo Fisher Scientific). For cultivation of human umbilical vein endothelial cells (HUVECs, Lonza, Basel, Switzerland), EBM™-2 Basal Medium (CC-3156, Lonza) supplemented with Hydrocortisone, hFGF-B, VEGF, R3-IGF-1, Ascorbic Acid, hEGF and GA-1000, all from EGM™-2 SingleQuots™ Supplement Pack (CC-4176, Lonza) abiding by manufacturer’s instructions as well as 10% (v/v) FBS was used. Human cardiac fibroblasts (HCF) were cultured in Fibroblast Growth Medium 3 (FGM-3), i.e. Fibroblast Basal Medium 3 (C-23230, PromoCell, Heidelberg, Germany) containing 10% (v/v) FBS, 1% (v/v) PenStrep as well as 1 ng/mL human recombinant basic FGF and 5 ng/mL Insulin (both from Growth Medium 3 SupplementPack, C-39350, PromoCell). HEK293FT and HUVECs were kept in T75 flasks (Sarstedt, Nümbrecht, Germany), HCFs in T150 flasks (TPP, Trasadingen, Switzerland), respectively, at 37 °C and 5% CO_2_ under regular exchanged to fresh culture medium and weekly passaging. For the latter, cells were washed with Dulbecco’s phosphate buffered saline (PBS, Invitrogen, Waltham, MA, USA) and 0.05% Trypsin-EDTA solution (Invitrogen) was applied for 5 min to 10 min achieving cellular detachment. Trypsinization was stopped with FBS containing medium and cells were pelleted for 5 min at 4 °C and 300°×g and resuspended in fresh culture medium for determination of cell concentration with Countess II (Thermo Fisher Scientific). For further cultivation 100000 HEK293FTs, 250000 HUVECs or 500000 HCFs per flask were seeded.

### Human cardiac fibroblasts

HCFs that were used in this study were either purchased from PromoCell (Lot-Nr. 436Z024.3, 450Z014.1, 452Z013.1) or isolated from tissue pieces from patients receiving a left ventricular assist device (application #9398_BO_K_2020 approved by Hanover Medical School (MHH) ethics committee, patients gave consent to use of material for research purposes, KFO311, MHH Register Herz-/Lungeninsuffizienz) kindly provided by the department of pathology (MHH). Tissue was washed for 5 min on ice with 1× Hanks’ balanced salt solution (HBSS, Gibco, Carlsbad, CA, USA) and cut into small pieces. Digestion was performed by applying 7 mL 1× HBSS containing 600 U/mL Collagenase II (Worthington Biochemical, Lakewood, NJ, USA) and 60 U/mL DNase I (AppliChem, Darmstadt, Germany) for 30 min at 37 °C with occasional up- and downpipetting. The suspension was stored on ice and remaining tissue pieces were digested with 5 mL of the aforementioned HBSS solution for 5 min to 30 min. The complete volume of the resulting suspension was cleared by filtering through a 100 µm filter (Miltenyi Biotec, Bergisch Gladbach, Germany) which was washed once with 1× HBSS afterwards. Centrifugation at 100 ×g for 2 min separated cardiomyocytes and the supernatant was centrifuged again at 300 ×g for 10 min to pellet remaining cells which were then cultivated for three days in FGM-3. Trypsinized cells were washed and resuspended in MACS buffer, i.e. BSA and autoMACS™ rinsing solution (Miltenyi Biotec) both 1:20 in ddH_2_O. One quarter of the of the cell suspension volume human anti-fibroblast Microbeads (Miltenyi Biotec) were added for 30 min at RT, light protected. Afterwards, two washing steps with 1 mL MACS buffer were performed before filtering the suspension through a 30 µm filter onto an MS column located in a suitable MACS separator (all Miltenyi Biotec), which had been rinsed with 500 µL MACS buffer before. After washing the column thrice with 500 µL MACS buffer for removal of unbound cells, Microbead bound cells were eluted by removing the column from the separator and flushing it with 1 mL MACS buffer upon pressure application. Obtained cells were washed once with MACS buffer and seeded in FGM-3 for further cultivation.

### NF-kB reporter Assay

HEK293FT were seeded into a 48-well plate (Nunclon™ Delta Surface, Thermo Fisher Scientific) at a density of 17500 cells per well. After 24 h, cells were transfected with pmiR-Report β-GAL plasmid expressing β-galactosidase for normalization and pSGN-luc plasmid expressing luciferase under an NF-kB promotor. For transfection, both plasmids were diluted in OptiMEM (51985, Thermo Fisher Scientific) at a concentration of 50 ng/mL and a Lipofectamine™ 2000 (11668. Thermo Fisher Scientific) solution (1:200 in OptiMEM) was prepared. After 5 min of incubation at RT, both solutions were mixed at equal volumes and left at RT for 20 min. Per well, 100 µL of the final mix were applied for 4 h. Afterwards, the transfection solution was removed and cell culture medium containing only 0.1% (v/v) FBS and either SGLT2i inhibitors (DAPA, S1548; EMPA, S8022; both Selleckchem, Planegg, Germany) or respective dimethyl sulfoxide (DMSO, A994, Carl Roth, Karlsruhe, Germany) control (Ctrl) alone or in combination with 300 ng/mL polyinosinic:polycytidylic acid (polyIC, P9582, Sigma-Aldrich, St. Louis, MO, USA) was added for 24 h. For harvesting, cells were incubated on ice for 10 min, detached by pipetting and pelleted by centrifugation at 300 ×g and 4 °C for 5 min. The pellet was resolved in 100 µL Cell Culture Lysis Reagent (Luciferase Assay System, E1500, Promega, Madison, WI, USA) on ice and cleared by centrifugation (8000 ×g, 5 min, 4 °C). For luciferase activity determination, 5 µL of the supernatant were diluted in 45 µL of Luciferase Assay Reagent (Luciferase Assay System, E1500, Promega) and measured at a Synergy HT reader (BioTek, Winooski, VT, USA). For normalization, β-galactosidase activity was determined using the β-Galactosidase Enzyme Assay System (E2000, Promega) according to manufacturer’s protocol. Absorbance was detected at a Synergy HT.

### Treatment of HUVECs and HCFs

If not stated otherwise, HUVECs were treated with 100 µM SGLT2i or respective Ctrl in EBM-2 containing 0.1% (v/v) FBS and 1% (v/v) PenStrep either with or without 300 ng/mL polyIC for 24 h before the start of the respective assay. HCFs were subjected to 100 µM SGLT2i or Ctrl in FGM-3 for 48 h prior to assay start. If stated, stimulation with transforming growth factor β (TGF-β, 240-B, R&D Systems, Minneapolis, MN, USA), dissolved in 0.1% (weight/volume [w/v]) bovine serum albumin (BSA, 810683, Sigma-Aldrich) in 4 mM HCl, was performed alongside the aforementioned drug application. In any case, the treatment was initiated 24 h to 48 h after the cells had been seeded and had reached sufficient confluence.

### Determination of mRNA expression levels

Cells were harvested in 1 mL QIAzol Lysis Reagent (Qiagen, Hilden, Germany) for 5 min at RT and 200 µL chloroform (Sigma-Aldrich) was added. After vigorous shaking, phases were separated by incubating at RT for 3 min and subsequent centrifugation at 12000 ×g and RT for 5 min. The aqueous phase was mixed with an equal volume of 2-propanol (Sigma-Aldrich) and incubated on ice for 10 min. RNA was precipitated by centrifuging the solution for 10 min at 12000 ×g and 4 °C. The RNA pellet was washed twice with 75% (v/v) Ethanol followed by a centrifugation for 10 min at 12000 ×g and 4 °C. The dried pellet was reconstituted in 20 µL RNase-free water and concentration as well as purity of RNA was measured at a Synergy HT. If needed, remaining cellular DNA was digested using 0.0164 Kunitz Units/µL DNase I in Buffer RDD (79254, Qiagen), RNA digestion was prevented by utilizing 0.533 U/µL RNasin^®^ ribonuclease inhibitor (N2511, Promega). After 30 min of incubation at 37 °C, 1.225 mM EDTA (15575020, Invitrogen) was added and the sample was heated to 65 °C for 5 min. Complementary DNA (cDNA) was generated using the Biozym cDNA Synthesis Kit (331470, Biozym, Hessisch Oldendorf, Germany) according to manufacturer’s protocol for Oligo-(dT) primers. Quantitative real time polymerase chain reaction (qPCR) was performed using the iQ™ SYBR® Green Supermix (170888, Bio-Rad, Hercules, CA, USA) according to manufacturer’s instructions. Prepared qPCR mixes of 10 µL contained respective primers (see **Supplemental Table S2**) at a concentration of 0.5 µM, 1:10000 ROX reference dye (from ABsolute Blue QPCR Mix, AB4136B, Thermo Fisher Scientific) and 1:400 Precision Blue™ (172555, Bio-Rad) as well as cDNA in a concentration ranging from 1 ng/µL to 4 ng/µL depending on target abundance. Reactions were run in 384-well plates (Bio-Rad) using a QuantStudio™ 7 Flex device and results were analyzed with QuantStudio™ Real-Time PCR software (both ABI, Waltham, MA, USA)

### Proteomics

HUVECs were seeded into 6-well plates at a density of 200000 cells per well, treated as described, and subsequently processed as described for Western Blot (see **Supplemental Methods**). Per sample, 18 µg of protein was prepared for gel loading by mixing with an appropriate volume of loading buffer containing DTT. Proteins were denatured for 5 min at 95 °C and cooled to RT before acrylamide (#1610140, Bio-Rad) was added to a final concentration of 4% (w/v) and the sample was incubated at RT for 30 min. Samples were loaded into wells of a Mini-PROTEAN^®^ TGX™ precast gradient (4% - 15%) gel (#4561083, Bio-Rad) and electrophoresis was run at 30 mA per gel for approximately 1 h. Proteins inside the gel were visualized by staining with FastGene^®^ Q-Stain (FG-QS1, Nippon Genetics, Düren, Germany) for 45 min. Before further processing, the stain was removed by washing the gel for 3 min in Millipore water. Proteins in gel lanes were digested with trypsin and separated with liquid chromatography prior to Orbitrap mass spectrometry analysis. Spectra were matched with Uniprot database (FDR < 0.01) resulting in 5646 identified proteins. For further analysis, only proteins identified in 83.3% of all samples were considered, leaving 3870 candidates. For normalization, respective median values were subtracted and missing values were imputed (normal distribution, reduction 1.8, width 0.3). Identified proteins with a peptide count less than 3 were excluded from further analyses. Principal component analysis was performed with the “prcomp” function of the “stats” package (version 3.6.2) for R with default settings except the “scales.”-argument was enabled. Statistically significantly regulated proteins were determined using Tukey’s HSD test (“TukeyHSD” function, R “stats” package version 3.6.2). For comparisons between selected treatment groups, proteins with absolute differences above 1 and adjusted p-values below 0.05 were subjected to overrepresentation analysis (ORA) for gene ontology (GO) terms using Enrichr (https://maayanlab.cloud/Enrichr/). For gene set enrichment analysis (GSEA), proteins were ranked based on intensity differences within the analyzed pair. The “gseGO” function of the “clusterProfiler” package (version 3.0.4) of “BiocManager” (version 3.17) for R was utilized applying the following settings: ont = “ALL”, keytype = “UNIPROT”, minGSSize = 2, maxGSSize = 800, pAdjustMethod = “fdr”. Default values were used for all other arguments.

### Seahorse XF Mito Stress Test

HCFs or HUVECs were seeded into an Agilent XF96 plate (Agilent, Santa Clara, CA, USA) which had previously been coated with gelatin (#G9382, Sigma-Aldrich, St. Louis, MO, USA) solution (0.1% weight/volume in water) for 30 min at 37 °C at a density of 10000 cells per well and treated as described, except treatment of HUVECs was applied in EGM-2. For the assay, cells were washed twice with assay medium, i.e. Seahorse XF DMEM medium, pH 7.4 (103575-100, Agilent) supplemented with 1 mM pyruvate, 2 mM glutamine, and 10 mM glucose. Finally, 180 µL of assay medium was left on the cells and the sensor cartridge was loaded as described in the manufacturer’s instructions to achieve final concentrations of 10 µM oligomycin and FCCP, and 5 µM rotenone/antimycin A. The Mito Stress Test protocol was run at a Seahorse XFe96 analyzer (Agilent). Data was processed using Wave Desktop (version 2.6.1, Agilent). Normalization of measurements was achieved by imaging and counting cell nuclei stained with 3.33 µg/mL Hoechst33342 (Thermo Fisher Scientific) for 10 min using a Cytation1 and respective Gen5 software (version 3.11, both BioTek).

### Measurement of reactive oxygen species content

We utilized the DCFDA / H2DCFDA - Cellular ROS Assay Kit (ab113851, Abcam, Cambridge, UK) to measure the accumulation of reactive oxygen species (ROS) inside living cells. For that, HUVECs and HCFs were seeded into 96-well plates (TPP) at a concentration of 20000 cells per well and treated as described. For stimulation of ROS production, 50 µM or 500 µM H_2_O_2_ was added to HUVECs or HCFs, respectively. For the assay, cells were washed once with PBS and subsequently stained with 20 µM DCFDA for 45 min. After an additional washing step, 100 µL 1× Assay Buffer was added per well and fluorescence intensity (*λ*_Excitation_ = 485 nm, *λ*_Emission_ = 535 nm) was measured at a Cytation1 for 6 h. For statistical comparison of measured groups, areas under curves (AUCs) were determined from resulting kinetics.

### Migration Assay

For the migration assay, HCFs and HUVECs, seeded at densities of 30000 cells and 25000 cells per well, respectively, in a 96-well plate coated with 0.1% gelatin solution, were stained with 5 μM Hoechst33342 (Thermo Fischer Scientific) in culture medium light-protected at 37 °C for 20 min. Afterwards, cell layers were scratched with a 20 µL pipette tip (Sarstedt) to create an artificial wound. Detached cells were removed by washing with PBS, and 100 µL of respective medium with treatments or controls were added. Cellular migration was monitored for 24 h at 37 °C and 5% CO_2_ using a Cytation1 in combination with a BioSpa8 device and respective Gen5 and BioSpa OnDemand (version 1.03, BioTek) softwares by acquiring images in the “DAPI” channel (*λ*_Excitation_ = 377 nm, *λ*_Emission_ = 447 nm). In this assay, treatment of HCFs was initiated as described and maintained throughout the wound closure process. HUVEC treatment was started immediately after wound infliction and applied in normal growth medium to minimize stress on the cells. Background in images acquired with the Cytation1 device was subtracted with the image processing module of the Gen5 software selecting an adequate rolling ball radius of approximately twice the average nuclei size. Relative areas covered (RAC) were determined with Fiji.

### Tube formation assay

To monitor tube formation of HUVECs, 96-wells were filled with 40 µL Corning^®^ Matrigel^®^ Basement Membrane Matrix, LDEV-free (354234, Corning, Corning, NY, USA). After solidification, 7500 cells per well were seeded on top of the Matrigel^®^-coating and treated with 100 µM SGLT2i or respective Ctrl in EGM-2. Images in “BrightField High Contrast” channel were acquired every 4 h with a Cytation1 / BioSpa8 device combination. For quantification, images recorded in different z-levels were combined with the “Z Projection” module of Gen5 applying the “Focus Stacking” method and converted to RGB files for analysis with the Fiji “Angiogenesis Analyzer” plugin ^26^.

### Endothelial to mesenchymal transition assay

Endothelial to mesenchymal transition (EndMT) medium was prepared by supplementing EBM-2 with 10% (v/v) FBS, 10 ng/mL TGF-β, 10 ng/mL IL1β (200-01B, PeproTech, Cranbury, NJ, USA) as well as hydrocortisone, ascorbic acid, and GA-1000 from EGM-2 supplement pack according to manufacturer’s instructions. HUVECs were seeded at a concentration of 12500 cells per well in 24-well plates (TPP) and after 24 h, medium was exchanged to EndMT medium to induce transition. Cells were fed with fresh EndMT medium every 2 days to 3 days and images to observe morphological changes were captured every day with Cytation1 / BioSpa8. Treatments were applied as depicted in the scheme in **Figure 3A** and cells were harvested for gene expression analyses as described.

### CFSE proliferation assay

Proliferation of HCFs was measured with carboxyfluorescein succinimidyl ester (CFSE) staining coupled to flow cytometry. First, mitotic activity of HCFs was halted by applying nocodazole (InSolution™ Nocodazole, Merck, Darmstadt, Germany) at a concentration of 100 µM for 16 h. Next, synchronized HCFs were harvested and stained with 5 µM CellTrace™ CFSE (C34570, Invitrogen) in PBS for 20 min at RT. Unbound CFSE was quenched by addition of five volumes of DMEM containing 10% (v/v) FBS for 5 min at 37 °C. Subsequently, stained cells were pelleted (300 ×g, 5 min), resuspended in FGM-3, and 30000 cells per well were seeded into a 24-well plate. After 24 h, treatments were applied for 72 h and cells were trypsinized as described before, washed with PBS, and resuspended in PBS supplemented with 1% (v/v) FBS and 2.5 mM EDTA. Fluorescence intensity of each cell was determined using a CytoFLEX S (Beckman Coulter, Brea, CA, USA). For comparisons among treatments, a reference threshold dividing the histogram of the viable cell population of the Ctrl / vehicle (veh) group at approximately 50% was set and applied to all groups per biological replicate.

### SiRNA transfection

Seeding of HUVECs and HCFs was done according to the descriptions for the respectively conducted experiments 24 h prior to transfection. SiRNAs (all Santa Cruz Biotechnology, Dallas, TX, USA) were diluted in OptiMEM to a concentration of 20 nM. Transfection reagent solutions were prepared by diluting Lipofectamine™ 2000 (for HUVECs) or Lipofectamine™ RNAiMAX (13778, Thermo Fisher Scientific) 1:125 in OptiMEM (for HCFs) and incubated for 5 min at RT. After mixing both solutions at equal volumes and 20 min incubation at RT, 50 µL per well of the final mix were pipetted onto the cells. After 4 h transfection mixes were removed from the cells and treatments were added as described before.

### BrdU cell proliferation ELISA

HUVECs and HCFs were seeded in 96-well plates coated with 0.1% (w/v) gelatin solution at a concentration of 5000 cells per well and treated as outlined. Cellular proliferation was assessed with the BrdU cell proliferation ELISA (11647229001, Roche, Basel, Switzerland). In brief, after the described treatment period, 1:1000 5-bromo-2’-deoxyuridine (BrdU) labeling reagent was added to the cells overnight. Labeled cells were fixed and the assay was performed according to manufacturer’s protocol. Optical density at 370 nm was measured and background signal at 490 nm subtracted 10 min after substrate solution addition using a Synergy HT reader.

### Statistics

If not otherwise indicated, plots show means and 95% confidence intervals (CIs) and individual points represent biological replicates described by *n*, which each consisted of multiple technical replicates themselves. R (version 4.2.3) was utilized for statistical analysis. Datasets were evaluated for normal distribution and homoscedasticity using Shapiro-Wilk and Levene test, respectively (“shapiro_test” and “levene_test” functions, both R “rstatix” package version 0.7.2). Dependent on the number of independent variables, 2- or 3-way analysis of variance (ANOVA) was applied (“aov” function, R “stats” package version 3.6.2). In case of subject matching, respective “within”- and “between”-factors were defined (“anova_test” function, R “rstatix” package version 0.7.2). *Post hoc*, groups were compared pairwise using estimated marginal means test with Dunnett’s method for *p*-value adjustment (“emmeans” function, R “emmeans” package version 1.8.6). In case of matching, paired t-test with Bonferroni correction was applied. If not otherwise mentioned, groups receiving SGLT2is were always compared with the respective Ctrl and *p*-values below 0.05 are indicated above the respective treatment group.

## Results

### SGLT2is regulate inflammatory response in HEK293FT cells and HUVECs

Clinical data specifically focusing on inflammatory signaling in patients receiving SGLT2is have shown that these drugs decrease expression and secretion of pro-inflammatory cytokines ^27–29^ and reduce inflammasome activation ^30^ through glucose-lowering and ketogenesis. However, it is still not entirely clear whether there is also a direct interference of SGLT2i with inflammatory processes in cardiovascular cells. Therefore, we first focused on the NF-kB axis, undoubtedly a central inflammatory agonist in cardiovascular disease progression ^31,32^.

Monitoring of endogenous NF-kB activity in HEK293FT cells was used to determine the direct impact of DAPA and EMPA inflammatory signaling (**Figure 1A**). For initial screening, three different drug concentrations were used. In contrast to 1 µM and 10 µM, luciferase activity was reduced upon treatment with 100 µM DAPA, while as for EMPA a trend was observable (*p* = 0.1524) after stimulation of inflammatory signaling with polyIC. Since the large plasmid size did not allow performing the assay in HUVECs, we analyzed the abundance of phosphorylated p65 (pP65) as well as inhibitor of kB kinase (IKK) alpha (**Supplemental Figure S1A**) and screened expression levels of NF-kB downstream targets, i.e. intercellular adhesion molecule (ICAM) 1 and IL6 (**Figure 1B**). Both IL6 and ICAM1 levels were significantly reduced by DAPA and ICAM1 also by EMPA after addition of the inflammatory trigger polyIC. However, inhibition of SGLT2 could be excluded as the underlying mode of action in either cell type as per lack of expression (**Supplemental Figure S1B**).

**Figure 1:**
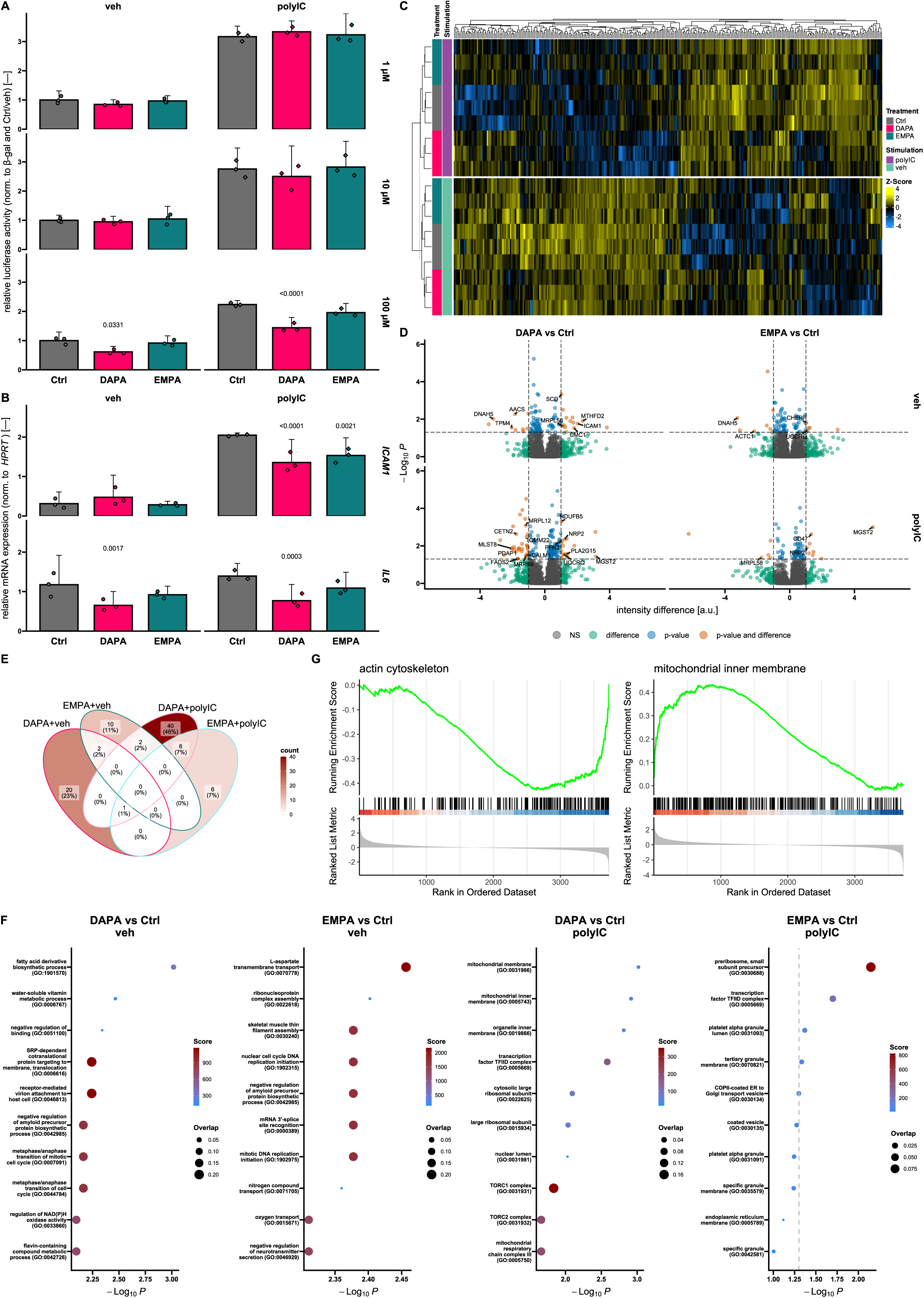
SGLT2is influence inflammatory signaling as well as mitochondrial parameters and actin cytoskeleton. **(A)** NF-kB dependent luciferase activity in human embryonal kidney 293FT reporter cells. Applied concentrations of dapagliflozin (DAPA) and empagliflozin (EMPA) are indicated on the right (*n* = 3). **(B)** Relative mRNA expression of NF-kB downstream targets intercellular adhesion molecule (*ICAM*) 1 (top) and interleukin (*IL*) 6 (bottom) in human umbilical vein endothelial cellls treated with 100 µM DAPA or EMPA or the respective control (Ctrl) (*n* = 3). **(C)** Heatmap showing by-protein Z-scores of intensities of proteins that are differentially regulated within the presented dataset (ǀintensity differenceǀ > 1 and adjusted *p*-value < 0.05). Hierarchical clustering was done based on Euclidean distance. Child and parent row dendrograms are separated by a dashed line (*n* = 3). **(D)** Volcano plots depicting dysregulated proteins by pairwise comparison (indicated above) in vehicle (veh) and polyinosinic:polycytidylic acid (polyIC) exposed HUVECs. **(E)** Venn-diagrams highlighting the overlap of all proteins that are regulated compared to the respective Ctrl among different groups investigated in proteomics analysis. **(F)** Analysis of pairwise dysregulated proteins (comparison indicated above) in veh and polyIC stimulated HUVECs for overrepresentation of gene ontology (GO) terms. Analysis was done with Enrichr. Dashed line indicates *p*-value cutoff. **(G)** Gene set enrichment analysis of proteins for comparison of DAPA (left) and EMPA (right) versus Ctrl in unstimulated HUVECs. Relevant terms are plotted, ranking was based on respective expression differences. a.u., arbitrary units; norm., normalized; β-gal, β-galactosidase; *HPRT*, hypoxanthine-guanine phosphoribosyltransferase.

### Impact of SGLT2i on the proteome of HUVECs

We investigated the proteome of ECs, a cell type controlling important roles during the course of inflammation ^33^, after exposure to SGLT2i and/or pro-inflammatory stimulus. In total, we identified 336 differentially expressed proteins in our dataset (**Figure 1C**) most of which were found by comparing veh and polyIC receiving cells, but differences between SGLT2i exposed and Ctrl cells were apparent (**Figure 1C, D** and **Supplemental Figure S2A**, **B**). In particular, DAPA regulated many proteins involved in cellular metabolism, mitochondrial processes and cytoskeleton. Interestingly, EMPA showed less profound effects on mitochondrial proteins as well as cytoskeletal parameters (**Figure 1D, F**). Deviant effects of the two investigated SGLT2is were identified as a substantial overlap in differentially expressed proteins between DAPA and EMPA was only observable when comparing upregulated candidates under polyIC stimulation (**Figure 1E** and **Supplemental Figure S2C**, **D**).

Additionally, we performed gene set enrichment analysis on selected terms ranking proteins according to absolute intensity differences for the respective comparison (**Figure 1G**). While an influence of DAPA on cytoskeletal components could be confirmed, EMPA interestingly showed an enrichment of mitochondrial inner membrane components. Under inflammatory conditions, DAPA led to a reduction of desmosome associated factors suggesting an influence on intercellular adhesion (**Supplemental Figure S2E**) even though, conversely, ECs are known to be devoid of desmosomal structures ^34,35^. We found an increase in glutathione biosynthesis after EMPA treatment in polyIC stimulated HUVECs suggesting a modulation of reactive oxygen homeostasis ^36^.

### DAPA but not EMPA interferes with HUVEC metabolism and mitochondrial respiration

Proteome analysis revealed a putative effect of SGLT2i on mitochondrial parameters. As no impact on intracellular mitochondrial structures by SGLT2is could be found (**Supplemental Figure S3A**), we performed Seahorse Mito Stress Test assay to identify whether particular processes of mitochondrial respiration in HUVECs are impacted. From the resulting oxygen consumption rate (OCR) profile, several parameters representing the respiratory state of the cells were calculated (**Figure 2A**). We observed a reduced maximal respiration combined with lower ATP production and spare respiratory capacity as well as non-mitochondrial oxygen consumption in unstimulated and stimulated, DAPA-treated HUVECs. Simultaneously, the extracellular acidification rate (ECAR) monitored during the Mito Stress Test assay was also diminished by DAPA (**Figure 2B** and **Supplemental Figure S3B**). At basal level, both of the investigated SGLT2i seemed to reduce ROS burden, while comparable to Seahorse data, this effect did not sustain stimulation of ROS production (**Figure 2C**).

**Figure 2:**
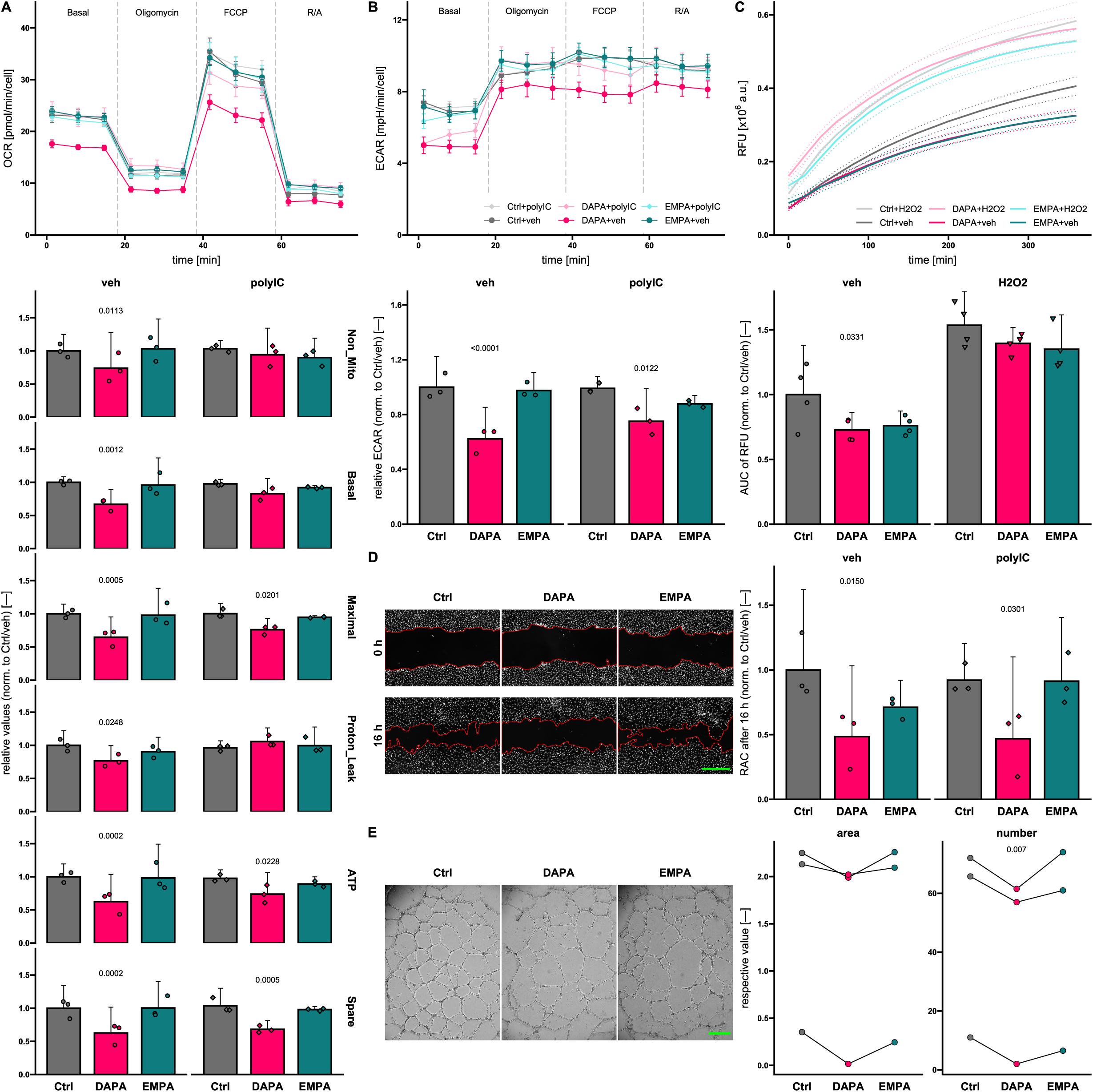
Sodium-glucose co-transporter 2 inhibitors influence HUVEC metabolism and migration. **(A)** Exemplary oxygen consumption rate (OCR) profile (top, 8 technical replicates) and values for mitochondrial respiration calculated based on OCR profiles (bottom, *n* = 3) of Seahorse Mito Stress test assays of human umbilical vein endothelial cells (HUVECs) treated as indicated in EGM-2. In upper graph, dashed lines separate measurement intervals between injections of different compounds (described above, “Basal“ means no compound added), means and standard errors are shown for measurements at each time point. FCCP, carbonyl cyanide-p-trifluoromethoxyphenylhydrazone; R/A, rotenone/antimycin A; Non_Mito, non-mitochondrial oxygen consumption; Basal, basal respiration; Maximal, maximal respiration; ATP, ATP production; Spare, spare respiratory capacity. **(B)** Representative extracellular acidification rate (ECAR) profile (top, 8 technical replicates) and ECAR values normalized (norm.) to Control (Ctrl) / vehicle (veh) group at the “Basal“ assay stage (bottom, *n* = 3). **(C)** Exemplary graph illustrating reactive oxygen species (ROS) levels in HUVECs treated as indicated assessed with H_2_DCFDA assay (top, 6 technical replicates). Means (solid lines) and standard errors (dashed lines) are shown. a.u., arbitrary units. Area under curve (AUC) of relative fluorescence units (RFU) curves (bottom, *n* = 4). **(D)** Relative areas covered (RAC) of the initial wound area in scratch assays of human umbilical vein endothelial cells (HUVECs) normalized to control (Ctrl) treated, vehicle (veh) receiving cells (right, *n* = 3). Representative “DAPI“ channel images of HUVECs (veh group) stained with Hoechst33342 are shown on the left. Wound borders at 0 h and 16 h are outlined in red. **(E)** Area and number of meshes formed by (treated) HUVECs on Matrigel after 24 h were evaluated by processing acquired z-projected bright field images (examples on the left) with the Angiogenesis Analyzer plugin for Fiji. Paired t-testing had to be applied due to high variations among the individual biological replicates (*n* = 3). Scale bar (green), 500 µm. DAPA, dapagliflozin; EMPA, empagliflozin; polyIC, polyinosinic:polycytidylic acid.

### DAPA diminishes HUVEC migratory and angiogenic activity

We performed high-throughput migration assays to elucidate the influence of SGLT2is on migration of ECs since proteomics results suggested altered cytoskeletal parameters (**Figure 2D**). Regardless of treatment with polyIC, DAPA reduced HUVEC migration significantly. As for EMPA however, a tendency towards reduced migratory capacity in unstimulated conditions can be assumed but was not statistically significant (*p* = 0.1771). To additionally analyze angiogenic characteristics, we seeded HUVECs on Matrigel^®^ to assess their tube formation capability (**Figure 2E**). While only a tendency towards mesh area reduction was visible, significantly fewer meshes were formed in total by HUVECs under exposure to DAPA. EMPA on the other hand did not influence HUVEC tube formation.

### Both investigated SGLT2is regulate mesenchymal transition of HUVECs

Endothelial to mesenchymal transition (EndMT) is frequently occurring during cardiac or vascular lesions enabling angiogenesis to reestablish nutrient supply to damaged tissue on the one hand but also contributing significantly to fibrotic tissue remodeling on the other hand ^37^. Mainly driven through TGF-β signaling ^38^, EndMT is characterized by a profound shift of the gene expression profile and functional phenotype of ECs which have recently been elaborated in great detail by Tombor *et al*. ^39^. Since DAPA and, to a lesser extent, EMPA seemed to exert influence on most EndMT characteristic parameters in our experimental settings, we performed *in vitro* EndMT assays with HUVECs and treated these cells with SGLT2is at different time points (**Figure 3A** and **Supplemental Figure S4A**). We analyzed a total of 14 genes representing different aspects of EndMT and observed morphological changes (**Figure 3B**, **C** and **Supplemental Figure S4B**, **C**). Even though HUVECs did not regain *PECAM1* expression through DAPA or EMPA treatment, the expression of the mesenchymal markers *TAGLN* and *CNN1* could be significantly suppressed. Interestingly, while this effect was generally visible for DAPA, EMPA treatment seemingly needed to be sustained for a longer period of time to achieve similar results. The same observations could be made for *COL1A1* expression. In line with previous results on inflammatory signaling, *IL6* overexpression during EndMT in this assay could also be partly repressed by SGLT2is. In line with the Seahorse data described earlier, *PFKP* was downregulated by EMPA (and DAPA, *p* = 0.0612) when applied with the start of the transition. Even though EndMT undergoing ECs show enhanced proliferation *in vivo* especially in the acute phase ^39,40^, markers for cell cycle arrest (*CDKN2A* and *CDKN1A*) were significantly elevated in transitioned cells *in vitro*. This phenotype could be consistently reversed by SGLT2i application.

**Figure 3:**
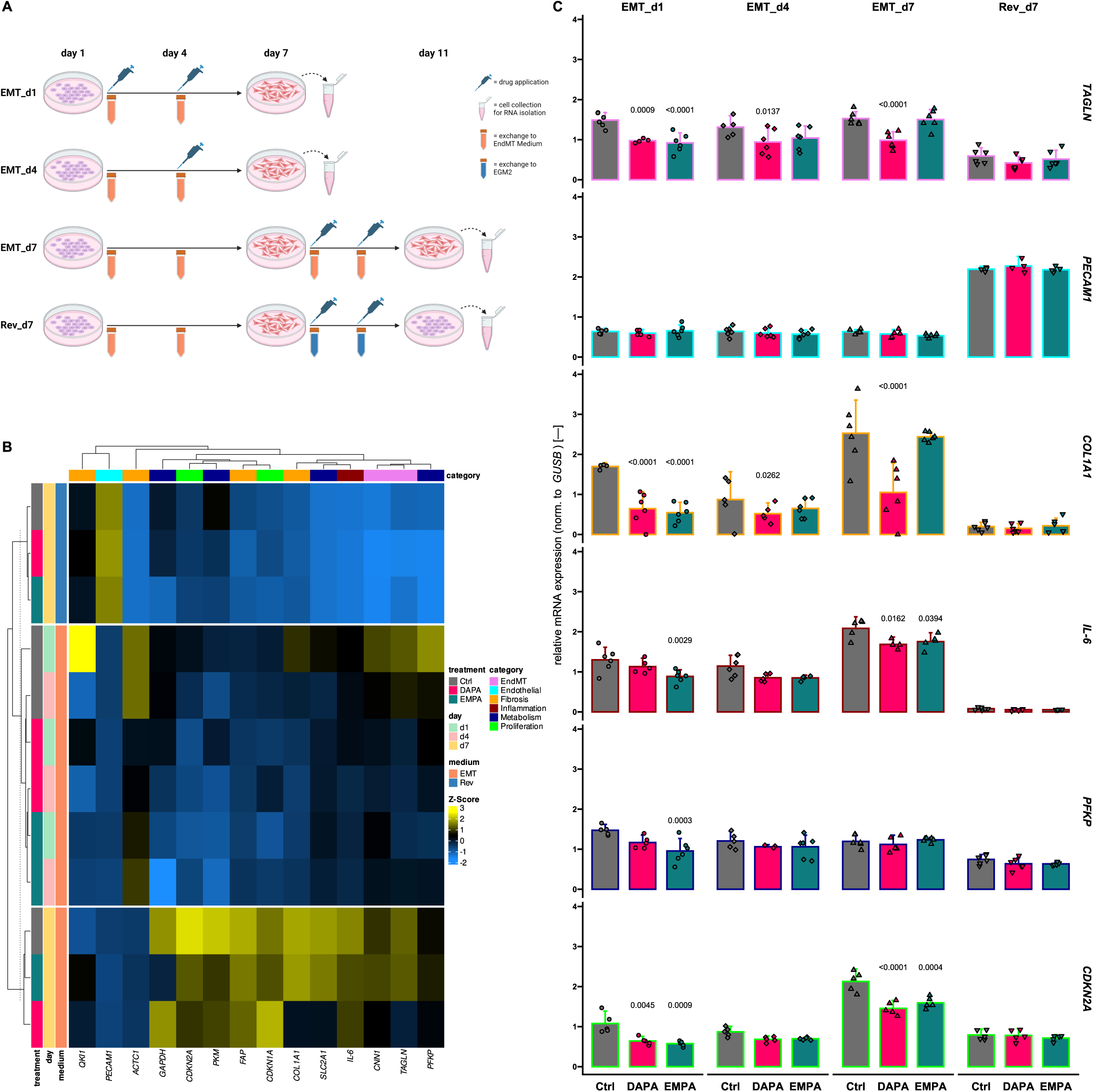
SGLT2i regulate EndMT. **(A)** Schematic representation of the experimental design to assess the influence of sodium-glucose co-transporter inhibitors (SGLT2is) on endothelial to mesenchymal transition (EndMT). Human umbilical vein endothelial cells (HUVECs) were seeded on day 0, 24 h prior to exchange of media. Groups were described as “EMT“, receiving EndMT medium from day 1 until the end of the experiment, or “Rev“, changing to EGM-2 after 6 days of EndMT induction. The second part of the name indicates the first day of treatment application. **(B)** Heatmap showing by-gene Z-scores of mRNA expression levels normalized (norm.) to β-glucuronidase (*GUSB*) of several genes from different categories assessed by quantitative real-time polymerase chain reaction. Hierarchical clustering was done based on Euclidean distance. Child and parent row dendrograms are separated by a dashed line (*n* ≥ 4). **(C)** Relative expression of representative genes for each assessed category. Bar outlines are colored according to the “category“ legend in **(B)**. DAPA, dapagliflozin; EMPA, empagliflozin; Ctrl, control; *TAGLN*, transgelin; *PECAM*, platelet endothelial cell adhesion molecule; *COL*, collagen; *IL*, interleukin; *PFKP*, phosphofructokinase, platelet; *CDKN*, cyclin-dependent kinase inhibitor.

### DAPA is anti-fibrotic via reduction of fibroblast migration and proliferation

We reported a strong effect of SGLT2is on EndMT, which is also arguably playing a pivotal role in fibrosis initiation and manifestation ^39,41,42^. Thus, we evaluated the impact of DAPA and EMPA on fibroblast biology using HCFs. In contrast to HUVECs, which are a pool of different donors, HCF lines were isolated from material of single patients. To enhance reproducibility and translational value, we conducted assays with multiple HCF lines including ones from tissue described as “healthy” (purchased from PromoCell) as well as self-isolated ones from patients suffering end-stage HF. The only significantly regulated OCR-based parameter in Seahorse Mito Stress test assay was ATP production that was suppressed by DAPA treatment (**Figure 4A**) while ECAR remained unaltered (**Figure 4B** and **Supplemental Figure S5A**). Consistent with the unaltered non-mitochondrial OCR, measurements of ROS accumulation showed no differences among compared (**Supplemental Figure S5B**). In congruence with the lowered ATP production, DAPA treated HCFs also showed decelerated proliferation (**Figure 4C** and **Supplemental Figure S5C**) as well as slower migration (**Figure 4D**). The effect was even more pronounced in pro-fibrotic conditions as treatment with TGF-β shows.

**Figure 4:**
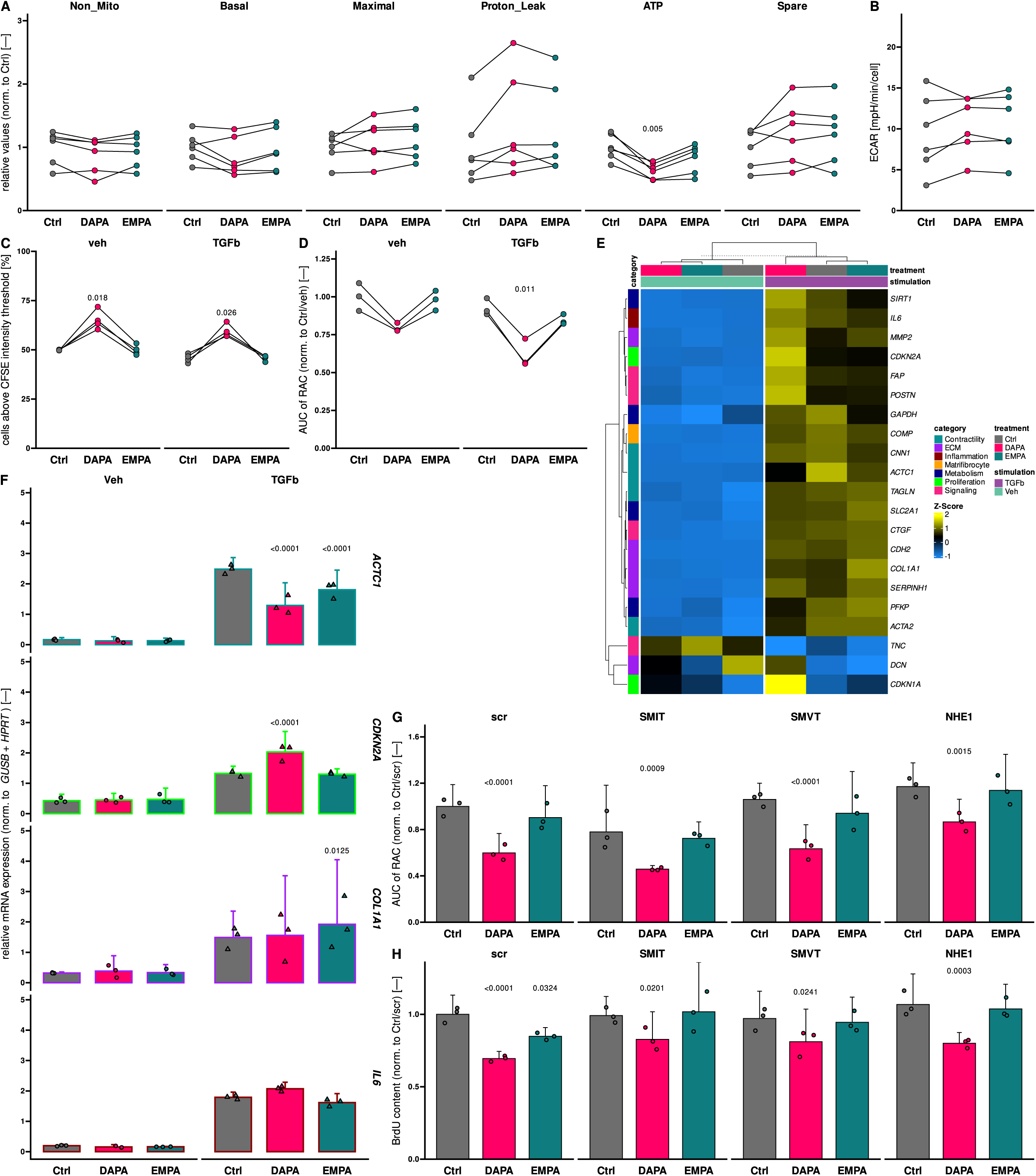
Dapagliflozin regulates fibroblast proliferation and migration. **(A)** Values for mitochondrial respiration calculated based on oxygen consumption rate profiles of primary human cardiac fibroblasts (HCFs) from different origins. Non_Mito, non-mitochondrial oxygen consumption; Basal, basal respiration; Maximal, maximal respiration; ATP, ATP production; Spare, spare respiratory capacity (*n* = 6). **(B)** Extracellular acidification rate values at the different Seahorse Mito Stress test assay “Basal“ stage. FCCP, carbonyl cyanide-p-trifluoromethoxyphenylhydrazone; R/A, rotenone/antimycin A (*n* = 6). **(C)** Percentage of carboxyfluorescein succinimidyl ester stained HCFs above the defined intensity threshold (*n* = 4). **(D)** Area under curve (AUC) of relative areas covered (RAC) of the initial wound area in scratch assays of HCFs over time normalized to control (Ctrl) treated, vehicle (veh) receiving cells (*n* = 3). **(E)** Heatmap showing by-gene Z-scores of mRNA expression levels in human cardiac fibroblasts (HCFs) receiving indicated treatments. Expression levels of several genes from different categories assessed by quantitative real-time polymerase chain reaction were normalized (norm.) to β-glucuronidase (*GUSB*) and hypoxanthine-guanine phosphoribosyltransferase (*HPRT*). Hierarchical clustering was done based on Euclidean distance. Child and parent column dendrograms are separated by a dashed line (*n* = 3). **(F)** Relative expression of representative genes for each assessed category. Bar outlines are colored according to the “category“ legend in **(E)**. **(G)** Area under curve (AUC) of relative areas covered (RAC) of the initial wound area in scratch assays of human umbilical vein endothelial cells over time normalized to control (Ctrl) treated cells transfected with scrambled siRNA (scr) (*n* = 3). **(H)** Relative content of 5-bromo-2’-deoxyuridine (BrdU) incorporated over 24 h into DNA of replicating HCFs under indicated conditions. Targets of siRNA transfections are indicated above (*n* = 3). DAPA, dapagliflozin; EMPA, empagliflozin; TGFb, transforming growth factor β; *ACTC1*, cardiac muscle alpha actin; *CDKN*, cyclin-dependent kinase inhibitor; *COL*, collagen; *IL*, interleukin; SMIT, sodium-myoinositol transporter 1; SMVT, sodium-multivitamin transporter; NHE, sodium-proton exchanger.

Since proliferation and migration are characteristic signs of fibroblast activation, we investigated whether the observed functional effects are rooted in interferences with fibroblast activation *per se*. We screened expression of a panel of 21 genes covering different aspects of fibroblast activation (**Figure 4E**). Interestingly, while contractility-tagged markers and those reflecting proliferative activity were regulated in stimulated HCFs, markers from other investigated categories were predominantly unchanged (**Figure 4F** and **Supplemental Figure S5D**).

### Effect of SGLT2is on HUVECs and HCFs are mediated through interference with SLC5A family and NHE1

To elucidate potential underlying modes of actions of the effects of DAPA and EMPA on HUVECs and HCFs, we investigated whether siRNA-mediated knockdown (KD) of putative candidates would interfere with the previously observed impacts. Employing the Swiss Target Prediction tool (http://www.swisstargetprediction.ch/) highlighted several members of the solute carrier (*SLC*) 5A family (**Supplemental Figure S6A**). Therefore, we analyzed the expression of *SLC5A* members and *SLC9A1* (encoding for NHE1), a known target of SGLT2is ^17–19^, in HUVECs and HCFs (**Supplemental Figure S1B**) and excluded all non-expressed candidates from further experiments. Upon KD of the respective candidate (**Supplemental Figure S6B**), migration and proliferation assays were performed. Migration of HUVECs seemed to be slightly reduced upon SMIT (encoded by *SLC5A3*) KD and tended to be increased through NHE1 silencing. However, none of the investigated candidates seemed to influence the effects of SGLT2is. In scr controls, we could observe a significant downregulation of BrdU incorporation in DAPA and EMPA treated HCFs as described before (**Figure 4H**). The effect size of DAPA was reduced upon KD of all the investigated candidates while the effect of EMPA was annulled.

## Discussion

In the present study, we unraveled detailed effects of DAPA and EMPA on ECs and CFs. We identified anti-inflammatory effects as well as modulation of cellular metabolism, tube formation and migration activity of ECs applying proteomics analysis and subsequent functional validation experiments. Interestingly, while targeting of the NF-kB axis underlay the reduced inflammatory response in HEK293FT cells, we could not identify NF-kB as the affected mediator in HUVECs. It had previously been reported that SGLT2is have anti-inflammatory impacts *in vivo* and *in vitro* ^43–45^. Gaspari *et al*. connected anti-inflammatory actions of DAPA after TNFα stimulation in HUVECs to NF-kB signaling through observation of lowered transcriptional expression of NF-kB ^46^. However, NF-kB transcription factor activity is acutely regulated by phosphorylation and ubiquitination of respective proteins ^47^ which we have presented to not being altered by SGLT2is in HUVECs. The reduction in expression of pro-inflammatory markers by DAPA and EMPA might be mediated through other signaling pathways such as AP-1 which has been highlighted as another important player in inflammatory EC activation ^33^ and is crucially reliant on maintenance of the redox state of a cysteine residue in the DNA-binding fraction ^48^. Glutathione synthesis pathway enrichment in our proteomics data and lowered ROS production in response to SGLT2is reflected by the H_2_DCFDA assay and in the non-mitochondrial oxygen consumption ^49^ suggest an improved redox balance thereby possibly protecting AP-1 from aberrant activation.

Mitochondrial associated factors were overrepresented among significantly regulated proteins by SGLT2is and again Seahorse data showed interference of DAPA but not EMPA with mitochondrial respiration. Interestingly, Secker *et al*. ^50^ observed decreased mitochondrial respiration and increased ECAR in kidney epithelial cells in response to canagliflozin, which interfered with complex I of the electron transport chain through which the authors explained the elevated nephrotoxicity of canagliflozin compared to DAPA or EMPA. However, in our case, toxic effects of DAPA or EMPA were excluded by performing WST-1 turnover and LDH activity measurements (**Supplemental Figure S7**). Apart from the acidification through CO_2_ production in the tricarboxylic acid cycle, the ECAR is driven by lactate production which occurs as a metabolite of pyruvate if the latter is not shuttled into mitochondria and processed to acetyl-CoA or oxaloacetate ^51^. Hence, the observed reduction in ECAR upon DAPA treatment indicates less lactate production and, under consideration of OCR measurements, insinuates an influence of DAPA on glycolytic metabolism in HUVECs *per se*. This would reflect clinical data showing a reduced utilization of glucose as energy source which, as initially introduced, is being speculated to mediate cardioprotective functions of SGLT2i to a certain degree ^12–15^.

Enhanced metabolism of glucose as well as migratory and angiogenic activity are characteristic features of activated ECs undergoing EndMT ^39^, a crucial player in tissue damage response disputed to influence adverse remodeling in the long run ^39,41,42^. It has been previously shown that DAPA interferes with mesenchymal activation of ECs and cardiac fibrosis in diabetic rats and HUVECs subjected to a high-glucose environment ^52^. Applying TGF-β and IL1β instead of glucose, we showed that DAPA and EMPA are able to inhibit excessive mesenchymal activation of ECs independent glucose handling. Considering reduced migratory behavior of HUVECs and HCFs as well as decelerated proliferation of HCFs subjected to DAPA and similar tendencies observed after EMPA application, we deduce that SGLT2is modulate EC activation in response to stress stimuli and alleviate fibrosis progression. Conversely, expression analyses of several genes representing different stages of fibrosis ^53^ could not confirm an overall reduction of fibroblasts activation. Rather, candidates specifically involved in proliferation (e.g. *CDKN2A*) and migration (e.g. *ACTC1*) were regulated as expected.

To elucidate the precise mechanism underlying the effects of DAPA and EMPA in ECs and HCFs, we focused on expressed putative targets. Most relevant for this study, NHE1 facilitates cellular migration through local alteration of intracellular pH stimulating elongation of actin filaments for cellular protrusion forming ^54,55^. In contrast to previous reports showing that inhibition of NHE1 reduces epithelial carcinoma cell migration ^56,57^, we could not detect such an effect in HUVECs. Interestingly though, the anti-proliferative actions of SGLT2is on HCFs were compromised by knockdown of NHE1. Similar results as for NHE1 in terms of fibroblast proliferation were observed under silencing of SMIT and SMVT. The latter is highly expressed in various aggressive cancers ^58^ and biotin, one of its substrates ^59^, has already been directly shown to influence proliferation of choriocarcinoma cells ^60^. Comparably, SMIT has been reported to be an important driver of acute myeloid leukemia cell proliferation ^61^ and pharmacological inhibition with phlorizin, the substance DAPA and EMPA are derived from, led to reduced proliferation and migration of lung cancer cells ^62^. Based on the described reduction in effect size of SGLT2is on HCF proliferation upon respective KD, we conclude that anti-fibrotic cardiovascular effects of SGLT2is ^63–65^ are mediated through their direct binding and inhibition of SMIT, SMVT, and NHE1.

Considering clinical data in which DAPA and EMPA appear similarly cardioprotective at comparable dosages, the question may be posed as to why these two SGLT2is exhibited somewhat deviating effects on HUVECs and HCFs in our *in vitro* assays. This could be partly explained by the differences in selectivity/affinity for targeted structures as it is already known for the primary target ^66^. Despite sometimes lacking statistical significance, DAPA and EMPA mostly showed parallel effects in our *in vitro* assays. In addition, results obtained from the EndMT setup, clearly outlining drug actions in dependence of application duration, arguably insinuated such affinity discrepancy between the investigated SGLT2is. In support of this notion, a clinical trial investigating ertugliflozin reported no statistical superiority of the investigational drug in terms of treatment of cardiovascular disease in a diabetic context over placebo control ^67^. Besides, impacts on other important cell types in the cardiovascular system might deviate thereby equalizing the resulting *in vivo* phenotypes.

## Conclusion

In this report, we investigated the influence of two SGLT2is, DAPA and EMPA, on ECs and CFs. While we could detect an influence of EMPA on inflammatory response, an even stronger one was observed for DAPA, which also altered energy metabolism in HUVECs and showed reduction of migration and proliferation in HCFs. Employing an *in vitro* EndMT assay further underlined anti-fibrotic effects of both SGLT2is. Mechanistically, we could prove that interference with NHE1, SMIT and SMVT underlies SGLT2i actions observed in these cell types. Based on the results we generated in this study, we propose that cardiovascular protective effects of SGLT2is in clinical trials are in part mediated through (1) suppression of pro-inflammatory factors by activated ECs, (2) inhibition of EndMT, and (3) reduction of fibroblast activation (**Figure 5**).

**Figure 5:**
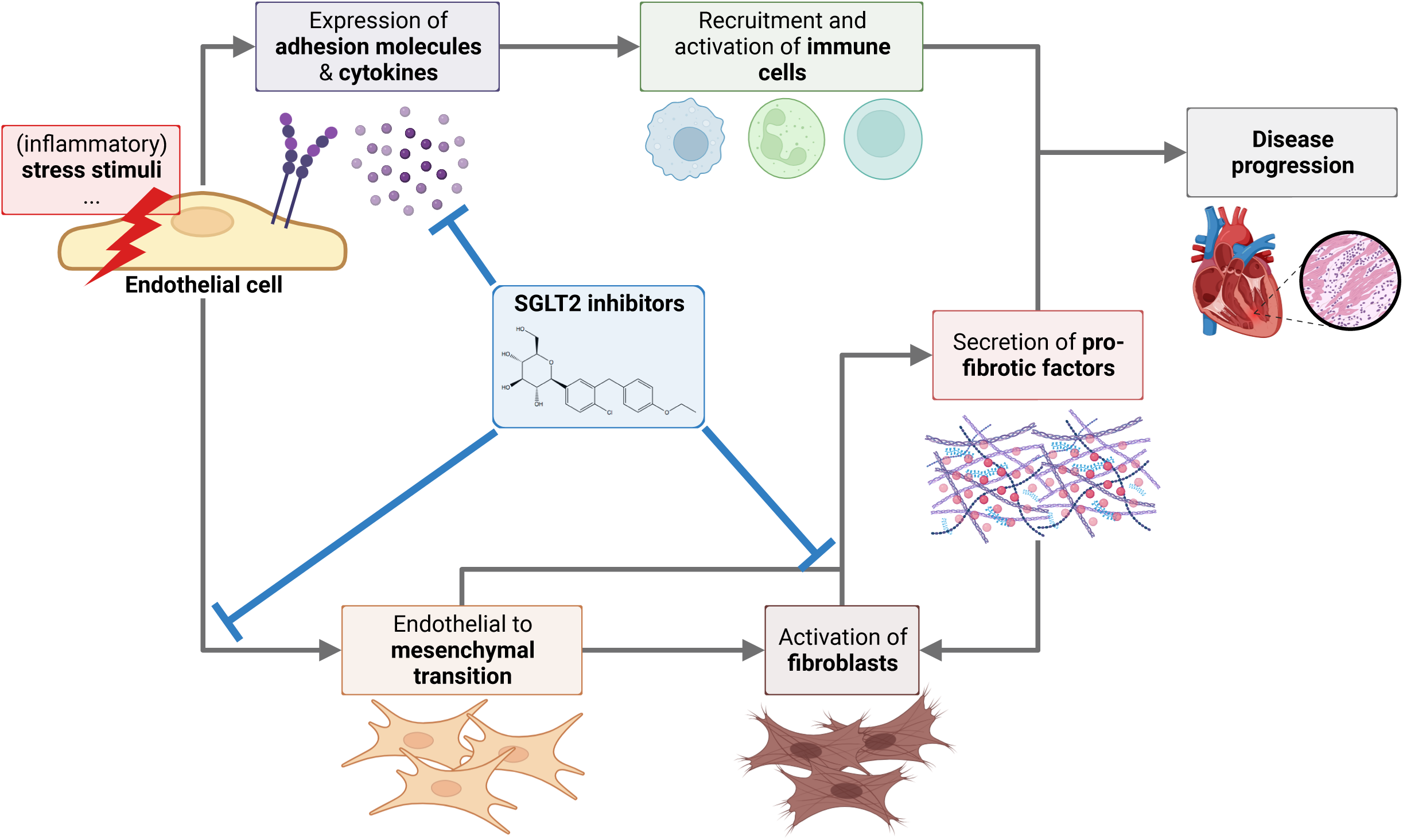
Overview of cardiovascular-protective effects of SGLT2is. SGLT2is regulate activation of endothelial cells after stress stimuli by interfering with expression of cytokines and adhesion molecules which are crucial for recruitment and activation of immune cells. Endothelial to mesenchymal transition is reduced by SGLT2is thereby altering pro-fibrotic signaling and fibroblast activation. A direct interference of SGLT2is with features of activated fibroblasts such as proliferation and migration reduces overall fibrotic load and adverse tissue remodeling ultimately hindering disease progression. Created in Biorender.com.

## Acknowledgements

K.S. and M.J. acknowledge support by the Hannover Biomedical Research School (HBRS).

## References

1. Ehrenkranz, J. R. L., Lewis, N. G., Ronald Kahn, C. & Roth, J. Phlorizin: a review. Diabetes. Metab. Res. Rev. 21, 31–38 (2005).

2. Uthman, L. et al. Direct Cardiac Actions of Sodium Glucose Cotransporter 2 Inhibitors Target Pathogenic Mechanisms Underlying Heart Failure in Diabetic Patients. Front. Physiol. 9, 1575 (2018).

3. Heerspink, H. J. L. et al. Dapagliflozin in Patients with Chronic Kidney Disease. N. Engl. J. Med. 383, 1436–1446 (2020).

4. Zinman, B. et al. Empagliflozin, Cardiovascular Outcomes, and Mortality in Type 2 Diabetes. N. Engl. J. Med. 373, 2117–2128 (2015).

5. Perkovic, V. et al. Canagliflozin and Renal Outcomes in Type 2 Diabetes and Nephropathy. N. Engl. J. Med. 380, 2295–2306 (2019).

6. Neal, B. et al. Canagliflozin and Cardiovascular and Renal Events in Type 2 Diabetes. N. Engl. J. Med. 377, 644–657 (2017).

7. Dhillon, S. Dapagliflozin: A Review in Type 2 Diabetes. Drugs 79, 1135–1146 (2019).

8. McMurray, J. J. V et al. Dapagliflozin in Patients with Heart Failure and Reduced Ejection Fraction. N. Engl. J. Med. 381, 1995–2008 (2019).

9. Packer, M. et al. Cardiovascular and Renal Outcomes with Empagliflozin in Heart Failure. N. Engl. J. Med. 383, 1413–1424 (2020).

10. Anker, S. D. et al. Empagliflozin in Heart Failure with a Preserved Ejection Fraction. N. Engl. J. Med. (2021) doi:10.1056/NEJMoa2107038.

11. Solomon, S. D. et al. Dapagliflozin in Heart Failure with Mildly Reduced or Preserved Ejection Fraction. N. Engl. J. Med. 387, 1089–1098 (2022).

12. Ferrannini, E., Mark, M. & Mayoux, E. CV Protection in the EMPA-REG OUTCOME Trial: A “Thrifty Substrate” Hypothesis. Diabetes Care 39, 1108–1114 (2016).

13. Mudaliar, S., Alloju, S. & Henry, R. R. Can a Shift in Fuel Energetics Explain the Beneficial Cardiorenal Outcomes in the EMPA-REG OUTCOME Study? A Unifying Hypothesis. Diabetes Care 39, 1115–1122 (2016).

14. Packer, M. SGLT2 Inhibitors Produce Cardiorenal Benefits by Promoting Adaptive Cellular Reprogramming to Induce a State of Fasting Mimicry: A Paradigm Shift in Understanding Their Mechanism of Action. Diabetes Care 43, 508–511 (2020).

15. Avogaro, A., Fadini, G. P. & Del Prato, S. Reinterpreting Cardiorenal Protection of Renal Sodium–Glucose Cotransporter 2 Inhibitors via Cellular Life History Programming. Diabetes Care 43, 501–507 (2020).

16. Dyck, J. R. B. et al. Cardiac mechanisms of the beneficial effects of SGLT2 inhibitors in heart failure: Evidence for potential off-target effects. J. Mol. Cell. Cardiol. 167, 17–31 (2022).

17. Uthman, L. et al. Class effects of SGLT2 inhibitors in mouse cardiomyocytes and hearts: inhibition of Na+/H+ exchanger, lowering of cytosolic Na+ and vasodilation. Diabetologia 61, 722–726 (2018).

18. Trum, M. et al. Empagliflozin inhibits Na+/H+ exchanger activity in human atrial cardiomyocytes. ESC Hear. Fail. 7, 4429–4437 (2020).

19. Trum, M., Riechel, J. & Wagner, S. Cardioprotection by SGLT2 Inhibitors—Does It All Come Down to Na+? International Journal of Molecular Sciences vol. 22 (2021).

20. Uthman, L. et al. Empagliflozin reduces oxidative stress through inhibition of the novel inflammation/NHE/[Na+]c/ROS-pathway in human endothelial cells. Biomed. Pharmacother. 146, 112515 (2022).

21. Li, X. et al. Sodium Glucose Co-Transporter 2 Inhibitors Ameliorate Endothelium Barrier Dysfunction Induced by Cyclic Stretch through Inhibition of Reactive Oxygen Species. International Journal of Molecular Sciences vol. 22 (2021).

22. Park, S.-H. et al. Angiotensin II-induced upregulation of SGLT1 and 2 contributes to human microparticle-stimulated endothelial senescence and dysfunction: protective effect of gliflozins. Cardiovasc. Diabetol. 20, 65 (2021).

23. Cappetta, D. et al. Amelioration of diastolic dysfunction by dapagliflozin in a non-diabetic model involves coronary endothelium. Pharmacol. Res. 157, 104781 (2020).

24. Ye, Y., Bajaj, M., Yang, H.-C., Perez-Polo, J. R. & Birnbaum, Y. SGLT-2 Inhibition with Dapagliflozin Reduces the Activation of the Nlrp3/ASC Inflammasome and Attenuates the Development of Diabetic Cardiomyopathy in Mice with Type 2 Diabetes. Further Augmentation of the Effects with Saxagliptin, a DPP4 Inhibitor. Cardiovasc. Drugs Ther. 31, 119–132 (2017).

25. Kang, S. et al. Direct Effects of Empagliflozin on Extracellular Matrix Remodelling in Human Cardiac Myofibroblasts: Novel Translational Clues to Explain EMPA-REG OUTCOME Results. Can. J. Cardiol. 36, 543–553 (2020).

26. Carpentier, G. et al. Angiogenesis Analyzer for ImageJ — A comparative morphometric analysis of “Endothelial Tube Formation Assay” and “Fibrin Bead Assay”. Sci. Rep. 10, 11568 (2020).

27. Garvey, W. T. et al. Effects of canagliflozin versus glimepiride on adipokines and inflammatory biomarkers in type 2 diabetes. Metabolism 85, 32–37 (2018).

28. La Grotta, R. et al. Anti-inflammatory effect of SGLT-2 inhibitors via uric acid and insulin. Cell. Mol. Life Sci. 79, 273 (2022).

29. Heerspink, H. J. L. et al. Canagliflozin reduces inflammation and fibrosis biomarkers: a potential mechanism of action for beneficial effects of SGLT2 inhibitors in diabetic kidney disease. Diabetologia 62, 1154–1166 (2019).

30. Kim, S. R. et al. SGLT2 inhibition modulates NLRP3 inflammasome activity via ketones and insulin in diabetes with cardiovascular disease. Nat. Commun. 11, 2127 (2020).

31. Fiordelisi, A., Iaccarino, G., Morisco, C., Coscioni, E. & Sorriento, D. NFkappaB is a Key Player in the Crosstalk between Inflammation and Cardiovascular Diseases. International Journal of Molecular Sciences vol. 20 (2019).

32. Gordon, J. W., Shaw, J. A. & Kirshenbaum, L. A. Multiple Facets of NF-κB in the Heart. Circ. Res. 108, 1122–1132 (2011).

33. Pober, J. S. & Sessa, W. C. Evolving functions of endothelial cells in inflammation. Nat. Rev. Immunol. 7, 803–815 (2007).

34. Bazzoni, G. & Dejana, E. Endothelial Cell-to-Cell Junctions: Molecular Organization and Role in Vascular Homeostasis. Physiol. Rev. 84, 869–901 (2004).

35. Vanslembrouck, B., Chen, J., Larabell, C. & van Hengel, J. Microscopic Visualization of Cell-Cell Adhesion Complexes at Micro and Nanoscale . Frontiers in Cell and Developmental Biology vol. 10 (2022).

36. Couto, N., Wood, J. & Barber, J. The role of glutathione reductase and related enzymes on cellular redox homoeostasis network. Free Radic. Biol. Med. 95, 27–42 (2016).

37. Welch-Reardon, K. M., Wu, N. & Hughes, C. C. W. A Role for Partial Endothelial– Mesenchymal Transitions in Angiogenesis? Arterioscler. Thromb. Vasc. Biol. 35, 303–308 (2015).

38. Dejana, E., Hirschi, K. K. & Simons, M. The molecular basis of endothelial cell plasticity. Nat. Commun. 8, 14361 (2017).

39. Tombor, L. S. et al. Single cell sequencing reveals endothelial plasticity with transient mesenchymal activation after myocardial infarction. Nat. Commun. 12, 681 (2021).

40. Virag, J. I. & Murry, C. E. Myofibroblast and Endothelial Cell Proliferation during Murine Myocardial Infarct Repair. Am. J. Pathol. 163, 2433–2440 (2003).

41. Zeisberg, E. M. et al. Endothelial-to-mesenchymal transition contributes to cardiac fibrosis. Nat. Med. 13, 952–961 (2007).

42. Piera-Velazquez, S. & Jimenez, S. A. Endothelial to Mesenchymal Transition: Role in Physiology and in the Pathogenesis of Human Diseases. Physiol. Rev. 99, 1281–1324 (2019).

43. Chen, S., Coronel, R., Hollmann, M. W., Weber, N. C. & Zuurbier, C. J. Direct cardiac effects of SGLT2 inhibitors. Cardiovasc. Diabetol. 21, 45 (2022).

44. Byrne, N. J., et al. Empagliflozin Blunts Worsening Cardiac Dysfunction Associated With Reduced NLRP3 (Nucleotide-Binding Domain-Like Receptor Protein 3) Inflammasome Activation in Heart Failure. Circ. Hear. Fail. 13, e006277 (2020).

45. Koyani, C. N. et al. Empagliflozin protects heart from inflammation and energy depletion via AMPK activation. Pharmacol. Res. 158, 104870 (2020).

46. Gaspari, T. et al. Dapagliflozin attenuates human vascular endothelial cell activation and induces vasorelaxation: A potential mechanism for inhibition of atherogenesis. Diabetes Vasc. Dis. Res. 15, 64–73 (2017).

47. Liu, T., Zhang, L., Joo, D. & Sun, S.-C. NF-κB signaling in inflammation. Signal Transduct. Target. Ther. 2, 17023 (2017).

48. Foppoli, C., Coccia, R. & Perluigi, M. Chapter 6 - Role of Oxidative Stress in Human Papillomavirus-Driven Cervical Carcinogenesis. in (ed. Preedy, V. B. T.-C.) 51–61 (Academic Press, 2014). 10.1016/B978-0-12-405205-5.00006-4.

49. Chacko, B. K. et al. The Bioenergetic Health Index: a new concept in mitochondrial translational research. Clin. Sci. 127, 367–373 (2014).

50. Secker, P. F. et al. Canagliflozin mediated dual inhibition of mitochondrial glutamate dehydrogenase and complex I: an off-target adverse effect. Cell Death Dis. 9, 226 (2018).

51. Mookerjee, S. A., Goncalves, R. L. S., Gerencser, A. A., Nicholls, D. G. & Brand, M. D. The contributions of respiration and glycolysis to extracellular acid production. Biochim. Biophys. Acta - Bioenerg. 1847, 171–181 (2015).

52. Tian, J. et al. Dapagliflozin alleviates cardiac fibrosis through suppressing EndMT and fibroblast activation via AMPKα/TGF-β/Smad signalling in type 2 diabetic rats. J. Cell. Mol. Med. 25, 7642–7659 (2021).

53. Fu, X. et al. Specialized fibroblast differentiated states underlie scar formation in the infarcted mouse heart. J. Clin. Invest. 128, 2127–2143 (2018).

54. Pollard, T. D. & Borisy, G. G. Cellular Motility Driven by Assembly and Disassembly of Actin Filaments. Cell 112, 453–465 (2003).

55. Stock, C. & Schwab, A. Role of the Na+/H+ exchanger NHE1 in cell migration. Acta Physiol. 187, 149–157 (2006).

56. Zhang, Y. et al. Polarized NHE1 and SWELL1 regulate migration direction, efficiency and metastasis. Nat. Commun. 13, 6128 (2022).

57. Lin, Y. et al. NHE1 mediates migration and invasion of HeLa cells via regulating the expression and localization of MT1-MMP. Cell Biochem. Funct. 30, 41–46 (2012).

58. Maiti, S. & Paira, P. Biotin conjugated organic molecules and proteins for cancer therapy: A review. Eur. J. Med. Chem. 145, 206–223 (2018).

59. Vadlapudi, A. D., Vadlapatla, R. K., Pal, D. & Mitra, A. K. Functional and molecular aspects of biotin uptake via SMVT in human corneal epithelial (HCEC) and retinal pigment epithelial (D407) cells. AAPS J. 14, 832–842 (2012).

60. Crisp, S. E. R. H. et al. Biotin supply affects rates of cellproliferation, biotinylation of carboxylases and histones, andexpression of the gene encoding the sodium-dependentmultivitamin transporter in JAr choriocarcinomacells. Eur. J. Nutr. 43, 23–31 (2004).

61. Wei, Y. et al. SLC5A3-Dependent Myo-inositol Auxotrophy in Acute Myeloid Leukemia. Cancer Discov. 12, 450–467 (2022).

62. Cui, Z. et al. The sodium/myo-inositol co-transporter SLC5A3 promotes non-small cell lung cancer cell growth. Cell Death Dis. 13, 569 (2022).

63. Li, C. et al. SGLT2 inhibition with empagliflozin attenuates myocardial oxidative stress and fibrosis in diabetic mice heart. Cardiovasc. Diabetol. 18, 15 (2019).

64. Chen, X. et al. Dapagliflozin Attenuates Myocardial Fibrosis by Inhibiting the TGF-β1/Smad Signaling Pathway in a Normoglycemic Rabbit Model of Chronic Heart Failure . Frontiers in Pharmacology vol. 13 (2022).

65. Yang, Z. et al. SGLT2 inhibitor dapagliflozin attenuates cardiac fibrosis and inflammation by reverting the HIF-2α signaling pathway in arrhythmogenic cardiomyopathy. FASEB J. 36, e22410 (2022).

66. Grempler, R. et al. Empagliflozin, a novel selective sodium glucose cotransporter-2 (SGLT-2) inhibitor: characterisation and comparison with other SGLT-2 inhibitors. *Diabetes*, Obes. Metab. 14, 83–90 (2012).

67. Cannon, C. P. et al. Cardiovascular Outcomes with Ertugliflozin in Type 2 Diabetes. N. Engl. J. Med. 383, 1425–1435 (2020).

